# Control of *C. elegans* growth arrest by stochastic, yet synchronized DAF-16/FOXO nuclear translocation pulses

**DOI:** 10.1101/2023.07.05.547674

**Authors:** Burak Demirbas, Olga Filina, Timo Louisse, Yvonne Goos, María Antonia Sánchez-Romero, María Olmedo, Jeroen van Zon

## Abstract

FOXO transcription factors are highly conserved effectors of insulin and insulin-like growth factor signaling, that are crucial for mounting responses to a broad range of stresses. Key signaling step is the stress-induced translocation of FOXO proteins to the nucleus, where they induce expression of stress response genes. Insulin signaling and FOXO proteins often control responses that impact the entire organism, such as growth or starvation-induced developmental arrest, but how body-wide coordination is achieved is poorly understood. Here, we leverage the small size of the nematode *C. elegans*, to quantify translocation dynamics of DAF-16, the sole *C. elegans* FOXO transcription factor, with single-cell resolution, yet in a body-wide manner. Surprisingly, when we exposed individual animals to constant levels of stress that cause larval developmental arrest, DAF-16/FOXO translocated between the nucleus and cytoplasm in stochastic pulses. Even though the occurrence of translocation pulses was random, they nevertheless exhibited striking synchronization between cells throughout the body. DAF-16/FOXO pulse dynamics were strongly linked to body-wide growth, with isolated translocation pulses causing transient reduction of growth and full growth arrest observed only when pulses were of sufficiently high frequency or duration. Finally, we observed translocation pulses of FOXO3A in mammalian cells under nutrient stress. The link between DAF-16/FOXO pulses and growth provides a rationale for their synchrony, as uniform proportions are only maintained when growth and, hence, pulse dynamics are tightly coordinated between all cells. Long-range synchronization of FOXO translocation dynamics might therefore be integral also to growth control in more complex animals.

## Introduction

FOXO transcription factors (TFs) are effectors of the insulin/insulin-like growth factor signaling (IIS) pathway that are essential for survival under adverse conditions [1], strongly linked to longevity [2,3] and implicated in diseases such as diabetes and cancer [4–6]. Surprisingly, in mammals only four distinct FOXO TFs are responsible for integrating signals for a highly diverse range of stresses, such as oxidative stress, nutrient depletion and DNA damage [7–9], to mount a response that is both tailored to the type of stress encountered and proportional to stress magnitude. The key step of IIS is signaling-induced translocation of FOXO TFs from the cytoplasm into the nucleus [10,11], where they induce stress-specific gene-expression programs.

So far, IIS and FOXO nuclear translocation are studied predominantly in cell culture, focusing on cellular responses to stress, such as cell cycle arrest and apoptosis [7–9]. However, a striking aspect of IIS is that it often elicits strong organismal responses. Both in *Drosophila* and mice, IIS and FOXO mutants show delayed body-wide growth [11–14], and IIS is responsible in *Drosophila* for adapting the rates of body-wide growth and development to changes in nutrients levels [15,16]. Moreover, in mammals FOXO TFs mediate the well-known metabolic response to changes in blood insulin level that is tightly coordinated between musculature, liver and adipose tissue [17]. However, FOXO nuclear translocation dynamics has barely been studied on the level of the entire organism.

Due to its small size and cell number, the nematode *Caenorhabditis elegans* is in principle ideally suited for studying cellular dynamics at the level of an entire body. Moreover, the IIS pathway is highly conserved from *C. elegans* to mammals (**Fig. 1A**), with a single *C. elegans* FOXO TF, DAF-16, controlling the response to stresses ranging from starvation, oxidative stress, osmotic shock to heat [18–21]. DAF-16/FOXO likely also controls organism-level decisions: for example, DAF-16/FOXO is required for developmental arrest of *C. elegans* larvae exposed to high stress [22–24]. However, the dynamics of nuclear DAF-16/FOXO translocation was never analyzed during developmental arrest, due to the difficulty of microscopy imaging in moving larvae. Instead, IIS and DAF-16/FOXO were only studied on the population level, by counting the fraction of animals with nuclear DAF-16/FOXO localization under different stresses. A puzzling result emerging from such studies is that even under constant, high stress, a substantial fraction (20-60%) of animals show no or little nuclear DAF-16/FOXO [21,25,26], hinting at a so far unresolved variability in DAF-16/FOXO response between individual larvae.

**Figure 1.**
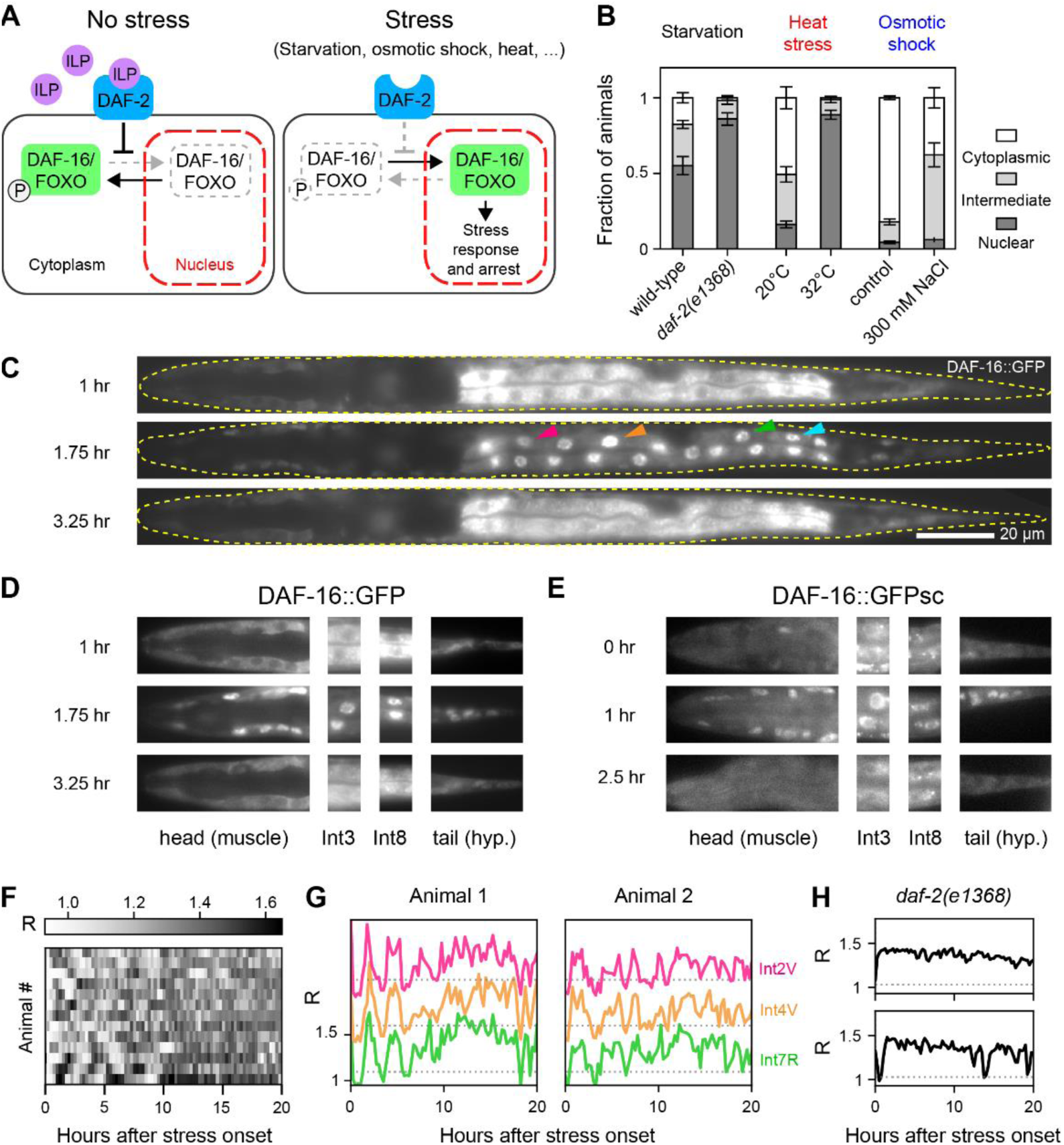
Synchronized, stochastic DAF-16/FOXO nuclear translocation pulses. **(A)** Schematic representation of the Insulin/insulin-like growth factor-1 signaling (IIS) pathway. **(B)** Fraction of L1 larvae with nuclear DAF-16::GFP localization for wild-type and *daf-2(e1368)* animals under starvation, and wild-type animals exposed to heat or osmotic shock. For each condition we performed 3 replicates of n>48 animals. Error bars are standard error of the mean (S.E.M.) over replicates. **(C)** Straightened fluorescence images of a starved L1 animal, carrying a DAF-16::GFP integrated transgene, showing alternating nuclear and cytoplasmic DAF-16 localization. Hours indicate time after hatching in microchambers without food. Dashed yellow line indicates animal’s outline. Scale bar: 20 µm. Arrowheads indicate intestinal cells Int2D/V (magenta), Int4D/V (orange), Int7L/R (green) and Int8L/R (cyan), used for single-cell quantification. **(D),(E)** DAF-16::GFP nuclear translocation dynamics in head muscle cells, intestinal cells, or tail hypodermal cells upon constant starvation stress. Both for DAF-16::GFP (D) and a single-copy, endogenously-tagged DAF-16::GFPsc strain (E), translocation occurred in pulses that were synchronized between the different tissues. Scale bar: 20 µm. Images in (C)-(E) were computationally straightened for clarity. **(F)** Heatmaps of DAF-16::GFP nuclear localization *R*, calculated as the ratio of DAF-16::GFP intensity in the nucleus and cytoplasm, over time. Rows represent individual, starved animals. Color intensity corresponds to *R*, as averaged over all intestinal cells. **(G)** DAF-16::GFP nuclear localization *R* measured for different intestinal cells, for two starved individuals. Despite variability between individuals, *R* is synchronized between cells. Tracks are averaged over two cells at the same antero-posterior (A-P) location, e.g. Int2D and Int2V, and are shifted along the y-axis for clarity. Dashed line is *R* =1.03, corresponding to 105% of the average DAF-16::GFP nuclear localization in non-stressed animals. **(H)** Nuclear localization *R*, averaged over all intestinal cells, for starved *daf-2(e1368)* mutants with constitutively low IIS activity.

Using a unique approach that allows time-lapse microscopy with single-cell resolution in freely-moving *C. elegans* larvae [27], we demonstrate here that, upon constant stress, DAF-16/FOXO enters and exits the nucleus in stochastic pulses of 1-2 hr duration, revealing that the perceived variability of nuclear DAF-16/FOXO levels between individuals in fact reflects stochastic pulse dynamics in time within each individual. Pulse dynamics also differed qualitatively between applied stresses, ranging from stochastic oscillations (starvation) to random pulses (osmotic shock) or a single pulse of fixed duration (heat shock). Surprisingly, we find that, despite their stochasticity, pulses are strikingly synchronized between DAF-16/FOXO expressing cells throughout the body: even when the exact time of a translocation pulse could not be predicted, it occurred simultaneously in all cells within <5 mins, pointing to a strong, but unknown synchronizing mechanism operating within individual animals. Moreover, we found that DAF-16/FOXO pulses were directly linked to body-wide growth arrest: individual DAF-16/FOXO translocation pulses often immediately (<0.5 hr) caused a transient halt in growth of body length. Moreover, full growth arrest occurred only when DAF-16/FOXO nuclear localization was of high frequency and/or long duration, suggesting that this arrest depends on DAF-16/FOXO translocation pulse integration. Finally, we show FOXO translocation pulses, which had so far not been observed in any model system, occur also in mammalian cells under nutrient deprivation, suggesting that they are a general feature of FOXO TFs.

The link between DAF-16/FOXO translocation pulses and *C. elegans* body growth provides a rationale for its synchronization, as it is essential that this stress response is executed uniformly throughout the entire organism. Strong body-wide synchrony might also be essential for stress responses unrelated to growth. Given our observation of translocation pulses also for mammalian FOXOs, we expect this novel, dynamic picture of synchronized nuclear translocation pulses to be important for understanding the key role of FOXO in mammalian biology, both at the cellular and organismal level.

## Results

### Synchronized, stochastic DAF-16/FOXO nuclear translocation pulses

To investigate organism-wide DAF-16/FOXO dynamics, we studied the *C. elegans* L1 arrest, a developmental arrest larvae enter when they encounter high stresses, including starvation, osmotic shock and heat, directly after hatching and that is under control of IIS [20,22–24,28,29] (**Fig. 1A**). Under unstressed conditions, high insulin-like peptide (ILP) levels activate the insulin receptor DAF-2, causing phosphorylation and cytoplasmic localization of DAF-16/FOXO [30–32]. Upon stress, low ILP levels result in DAF-2 inactivation, DAF-16/FOXO dephosphorylation and its subsequent translocation into in the nucleus, where it induces stress response genes [33–35] and regulators of cell proliferation [22], ultimately leading to developmental arrest.

We visualized DAF-16/FOXO nuclear localization using a transgenic line expressing a functional DAF-16::GFP fusion protein that tags the DAF-16a/b isoforms [36] and is a standard reporter for studying DAF-16/FOXO activation [23,37–40]. When we examined a population of starved L1 larvae 12 hours after hatching, DAF-16::GFP was nuclear in most animals, as expected, yet 20-40% of animals instead showed cytoplasmic localization (**Fig. 1B**), consistent with previous observations [21,25,26]. Similar variability was seen for 300 mM osmotic shock, while even for conditions that elicited a strong response (32°C heat shock or *daf-2(e1368)* mutants under starvation) still 5-10% of animals displayed cytoplasmic localization.

So far, this variability in DAF-16::GFP nuclear localization has only been studied on the population level, leaving open its activation dynamics in individual animals. We therefore performed time-lapse microscopy on animals hatched from eggs placed in 0.2x0.2 mm hydrogel microchambers [27] without food, allowing us to examine starvation-induced DAF-16/FOXO translocation dynamics in individual larvae (**SI Fig. 1A**). Upon hatching in absence of nutrients, DAF-16::GFP translocated from the cytoplasm to the nucleus within ∼2 hr in most animals, as expected. Surprisingly, DAF-16::GFP typically moved back into the cytoplasm after ∼1 hr (**Fig. 1C**) and continued to shuttle between the nucleus and the cytoplasm in pulses with ∼2 hr duration (**SI Movie 1**), even as starvation conditions were constant. Strikingly, even though timing and duration of DAF-16::GFP translocation pulses varied in time, DAF-16::GFP translocation was highly synchronized between *daf-16*-expressing cells throughout the body (**Fig. 1D, SI Movie 1**). We observed similar synchronized nuclear translocation pulses in a strain carrying a DAF-16::GFP endogenous CRISPR/Cas9 knock-in fusion (DAF-16::GFPsc) [41] (**Fig. 1E**), confirming that these translocation pulses were not an artefact of the DAF-16::GFP transgenic line. Because DAF-16::GFPsc fluorescence was much dimmer than DAF-16::GFP, we used DAF-16::GFP for quantifying pulse dynamics, even though its relative brightness indicated that it likely overexpresses DAF-16/FOXO.

We quantified DAF-16/FOXO nuclear translocation dynamics by measuring the ratio *R* between nuclear and cytoplasmic fluorescence (**SI Fig. 1B**, see **Methods** for details). We focused on intestinal cells, that form a stereotypical array along the antero-posterior (A-P) axis spanning approximately half the body length of L1 larvae. Most animals displayed initial oscillation-like translocation pulses (**Fig. 1F,G**), followed by more persistent DAF-16::GFP nuclear localization (*R*>1) as stress persisted, although brief pulses of translocation to the cytoplasm remained visible even at this stage. In general, *R* displayed clear variability between individual animals, in terms of both frequency and number of translocation pulses, underlining their stochastic nature. Yet, despite this individual variability, *R* dynamics always showed strong similarity between intestinal cells at markedly different positions along the A-P axis (**Fig. 1G**), revealing a degree of synchrony that was unexpected for such an intrinsically variable process.

We observed no DAF-16::GFP translocation pulses when eggs hatched in microchambers filled with plentiful food (**SI Fig. 1C**), indicating that they are a specific response to starvation. In *daf-2(e1368)* animals, that carry a mutation in the ligand binding domain of the DAF-2 insulin receptor [42] and therefore displayed a higher fraction of animals with nuclear DAF-16 in population-level measurements (**Fig. 1B**), we found that *R* dynamics was also changed: upon hatching without food, DAF-16::GFP was localized primarily in the nucleus and exhibited infrequent ∼1 hr pulses of translocation into the cytoplasm (**Fig. 1H)**, indicating that DAF-16::GFP pulse dynamics was controlled by IIS. Overall, these measurements reveal that the animal-to-animal variability in DAF-16::GFP nuclear localization under constant stress, that was previously observed on the population level, in fact reflects stochastic DAF-16::GFP translocation pulses occurring with high internal synchrony within each individual.

### Synchrony of DAF-16/FOXO translocation pulses is independent of stress type

We next asked if similar DAF-16::GFP nuclear translocation pulses were also seen for other types of stress. We exposed L1 larvae hatched in microchambers filled with food either to heat or osmotic shock, which are both known to induce DAF-16 nuclear translocation [23,36]. For osmotic shock, larvae were hatched in microchambers soaked in 200 or 300 mM NaCl, while for heat shock, larvae were shifted from 20°C to 32°C within 5 hrs after hatching (**SI Fig. 2A-D**, see **Methods** for details). Interestingly, both stress treatments resulted in DAF-16::GFP translocation pulses but with qualitative differences in dynamics, as will be examined in more detail further below. In particular, compared to starvation (**Fig. 1F,G**), translocation pulses under osmotic shock appeared more variable, both in time and between individuals (**Fig. 2A**), while heat shock induced a single translocation pulse that varied much less between individuals (**SI Fig. 2E**). Nevertheless, for both stresses the dynamics of *R* was still highly synchronized between intestinal cells. This was particularly striking for animals under 200 mM NaCl osmotic shock, where translocation pulses were often infrequent and of short duration, yet simultaneous in all cells examined.

**Figure 2.**
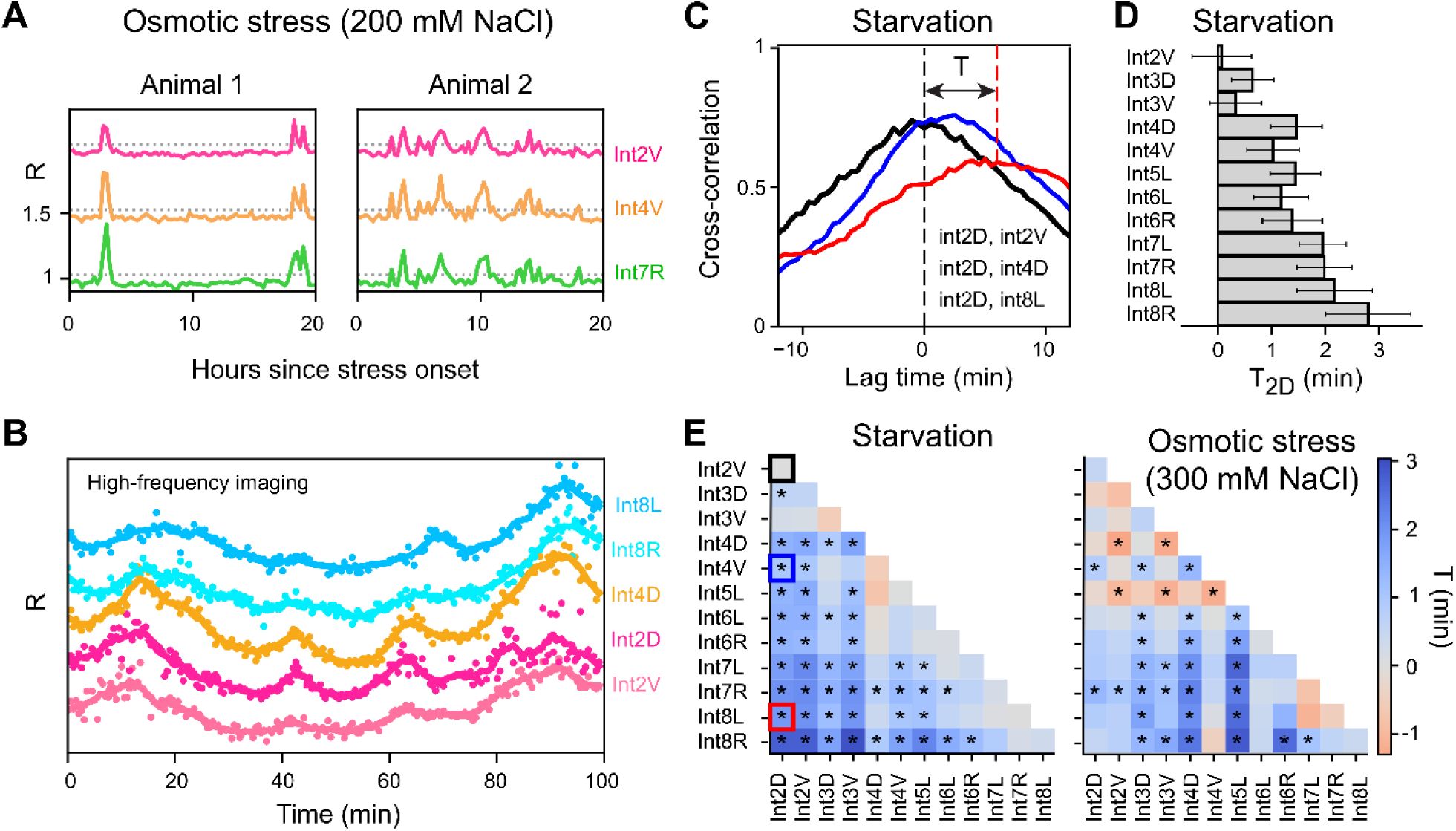
Synchrony of DAF-16/FOXO translocation dynamics in intestinal cells. **(A)** Synchronized DAF-16::GFP nuclear localization *R* in intestinal cells of animals exposed to 200 mM NaCl osmotic shock. **(B)** DAF-16::GFP nuclear localization *R* in intestinal cells of a starved L1 animal, imaged at 30-second time resolution (markers). Lines are data smoothed with a Savotsky-Golay (S-G) filter. Individual tracks are shifted along the y-axis for clarity. **(C)** Cross-correlation of *R* for int2D and intestinal cell pairs at the same (int2V) or increasing A-P distance (int4D and int8L, respectively). Data is for n=4 animals. The lag time T at which the cross-correlation peaks, corresponds to the average delay in *R* between cells. **(D)** Measured delay T2D between Int2D and all other intestinal cells. Error bars are 95% confidence intervals of the peak centroid distributions obtained from Monte-Carlo simulations (*n*=4 animals). **(E)** Heatmap showing the delay T between all pairs of intestinal cells for starvation (left, *n*=4 animals) and osmotic shock (right, *n*=4). Squares denoted by * indicate a delay that is significantly different from 0 (*T*=0 lies outside 95% confidence interval). Outlined squares correspond to the curves in (C).

To put an upper bound on the degree of synchrony, we imaged DAF-16::GFP pulses with high, 30 s time resolution in animals under starvation stress and quantified *R* in the intestinal cells Int2-Int8, excluding the four Int1 and two Int9 cells where DAF-16::GFP was difficult to resolve in single cells (**Fig. 2B**). At this increased time resolution, we could observe gradual nuclear translocation of DAF-16::GFP over a ∼20 min time interval. Moreover, we found that *R* displayed a shift in time between different cells, although the magnitude of the shift was small compared to the timescale of translocation. To quantify the shift, we cross-correlated *R* between each pair of cells (**Fig. 2C**). Here, the lag time at which the cross-correlation peaked represented an estimate of the delay in pulses between the two cells, with the error in the delay estimated through Monte-Carlo simulations (**SI Fig. 3A-D**).

We found that cells in close physical proximity within the body, such as Int2V and Int2D that have the same A-P position, showed no clear delay (**Fig. 2C-E**). However, we found that the cross-correlation showed a peak at positive lag time when we compared anterior cells to more posterior cells (**Fig. 2C**), with the magnitude of the delay increasing with the distance between the cells (**Fig. 2D,E**). Notably, the sign of the delay indicated an anterior-to-posterior order in DAF-16::GFP pulse dynamics, that was consistent between different animals, with pulse dynamics initiating in the Int2 cells ∼3 min before the Int8 cells. When analysing high time resolution data obtained for osmotic shock, we found an equally strong synchronization, with delays between cells of <3 min (**Fig. 2E**). Intriguingly, the nature of the delay was more complex than the simple anterior-to-posterior order seen for starvation, and instead reflected a partial dorsal-ventral ordering. For example, the anterior but ventral cells Int2V, Int3V and Int4V had pulse dynamics delayed compared to the more posterior but dorsal cells Int4D and Int5L (**SI Fig. 3E, F**). Overall, this suggests that the synchronizing mechanism depends at least partially on stress type.

### DAF-16/FOXO translocation dynamics encodes both stress type and magnitude

We then more systematically examined the differences in DAF-16::GFP translocation dynamics between different stresses. Given the observed synchrony between different intestinal cells, we here only compared *R* dynamics quantified over all intestinal cells. Differences in DAF-16::GFP translocation dynamics were already apparent when we compared population-averaged translocation dynamics, 〈*R*〉, between stresses (**Fig. 3A**): for 32°C heat shock, 〈*R*〉 exhibited a single translocation peak ∼1.5 hrs after stress onset, consistent with our observation of a single, stereotypical translocation pulse in individuals (**SI Fig. 2E**), followed by a more gradual increase in nuclear localization as stress persisted. A similar immediate peak in 〈*R*〉 was visible upon starvation, but here followed by additional peaks, consistent with our earlier observation of more regular, oscillation-like translocation pulses in individual animals (**Fig. 1F-G**). In contrast, no clear peaks in 〈*R*〉 were visible for 300 mM NaCl osmotic shock, as would in principle be expected for the random translocation pulses we observed in individuals (**Fig. 2A, 3C**).

**Figure 3.**
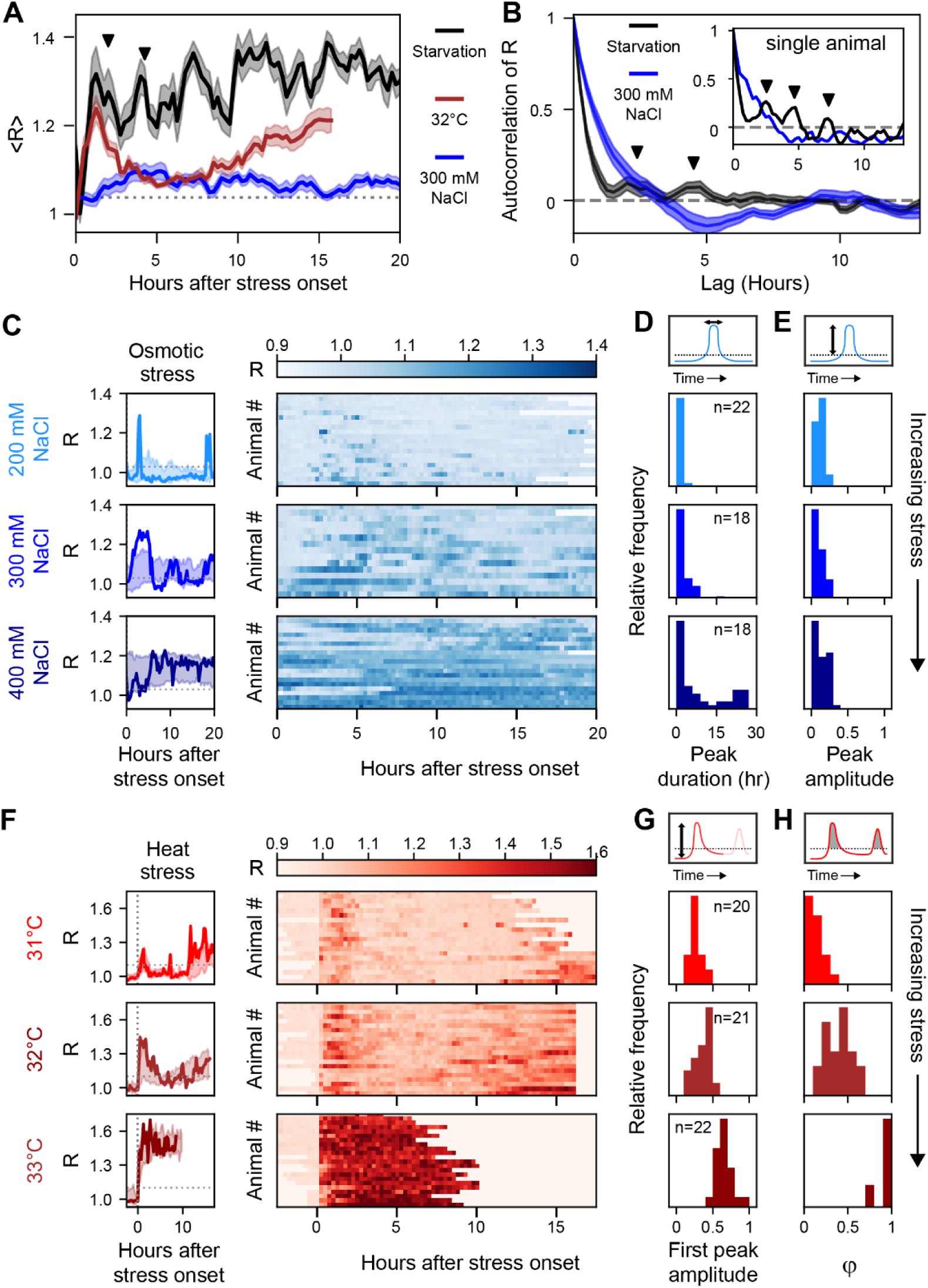
DAF-16/FOXO nuclear translocation dynamics depends on stress type and magnitude. **(A)** Population-averaged nuclear localization *R* for starvation (black, n=13), 300 mM osmotic shock (blue, n=18) and 32°C heat shock (red, n=21). Error bars are S.E.M. Arrowsheads indicate the population-average oscillation period of DAF-16::GFP translocated as determined in (B). **(B)** Population-averaged autocorrelation of *R* for starvation (black, n=13) and 300 mM osmotic shock (blue, n=18). Inset shows autocorrelation for an individual animal. Error bars are S.E.M. Peaks in autocorrelation (indicated by arrows) correspond to oscillatory DAF-16::GFP translocation dynamics. **(C)** Individual tracks (left) and population view heatmaps (right) of *R* for increasing osmotic shock. Individual tracks (left) are compared with the standard deviation for the population (shaded regions). **(D)**,**(E)** Distribution of peak duration (D) and peak amplitude (E) of DAF-16::GFP nuclear translocation pulses for osmotic shock. Increasing osmotic shock leads to more prolonged pulses without affecting amplitude. **(F)** Same as (C), but for increasing heat shock. For 33°C heat shock, observation time is limited to ∼10 h due to heat-induced shrinkage of the hydrogel microchambers. **(G)**,**(H)** Distribution of amplitude of the first DAF-16::GFP translocation peak (G) and fraction of time *ϕ* that DAF-16::GFP is strongly nuclear (H) for increasing heat shock.

To further differentiate *R* dynamics between starvation and osmotic shock, we determined the autocorrelation of *R* (**Fig. 3B**). Indeed, for starvation the autocorrelation in individual animals was oscillatory with exponentially decreasing envelope, a signature of stochastic oscillations. The population-averaged autocorrelation decayed more rapidly, showing only two peaks, at 2 and 4.25 hr, due to variation in pulse period and number of pulses between individuals. In contrast, the autocorrelation of *R* for osmotic shock showed simple exponential decay, both in individuals and averaged over all animals (**Fig. 3B**), consistent with random pulse dynamics.

Inspecting *R* dynamics in individuals highlighted the increased variability in number and duration of translocation pulses for osmotic shock, compared to starvation (**Fig. 1F, 3C**). For heat shock, we found that, even though the nature of the induced translocation dynamics was more stereotypical than for starvation or osmotic shock, substantial individual variability still remained. First, 6/21 animals failed to display a DAF-16::GFP pulse directly after shifting to 32°C. In addition, the gradual increase in 〈*R*〉 after the initial peak did not reflect a stereotypical increase in DAF-16::GFP nuclear localization in all individuals, but rather an increasing frequency and duration of pulses of increased nuclear localization that otherwise occurred stochastically within each individual.

In contrast to starvation, for osmotic and heat shock we could vary stress magnitude and examine the impact on DAF-16::GFP pulse dynamics. For osmotic shock, we hatched larvae in microchambers soaked with salt concentrations increasing from 200 to 400 mM NaCl. For 200 mM NaCl, we found that DAF-16::GFP displayed infrequent and short (<1 hr) pulses of nuclear localization, while increasing NaCl levels led to a proportional increase in the fraction of time DAF-16::GFP was nuclearly localized (**Fig. 3C**). This was due mainly to an increase in pulse duration (**Fig. 3D**), rather than pulse frequency (**SI Fig. 4A,B)**, and occurred without any concomitant increase in pulse amplitude (**Fig. 3E**), i.e. the degree of nuclear enrichment.

For heat shock, we shifted hatched L1 larvae from 20°C to temperatures ranging from 31°C to 33°C. DAF-16::GFP dynamics displayed similar characteristics both for 31°C and 32°C heat shocks: most animals displayed a single transient pulse of strong nuclear translocation directly after the shift to high temperature, followed by dynamic pulse-like changes in nuclear DAF-16::level (**Fig. 3F**). However, both the amplitude and duration of the initial translocation pulse increased with higher temperature (**Fig. 3G**, **SI Fig. 4C)**. In addition, the nuclear localization pulses that followed the initial pulse increased with temperature both in frequency and duration, resulting in a higher fraction of time ϕ that DAF-16::GFP was strongly nuclear (*R* >1.05, **Fig. 3H**). Following a 33°C heat shock, in contrast, we no longer observed a transient pulse, but found that DAF-16::GFP, after translocating, remained highly enriched in the nucleus for the duration of the experiment (**Fig. 3F**). In conclusion, for both osmotic and heat shock, higher stress magnitude was apparent through the increased fraction of time that DAF-16::GFP was nuclear.

Overall, this showed that DAF-16::GFP nuclear translocation dynamics was not only distinct for different types of stress (starvation, osmotic shock or heat shock), but also informed on stress magnitude. As DAF-16/FOXO mediates stress response primarily by binding to promoters of target genes and upregulating their expression [34,35,43], which crucially depends on DAF-16/FOXO nuclear localization, these observed differences in DAF-16/FOXO translocation dynamics thus likely impact stress response gene expression, in a manner that depends on stress type and magnitude.

### DAF-16/FOXO pulses are causally linked to arrest of body growth

We then examined possible functional roles of DAF-16/FOXO translocation pulses, focusing on the most prominent feature of L1 arrest, namely the arrest of body growth [22,28]. IIS and DAF-16/FOXO have a well-established role in inducing cell cycle arrest during starvation [22,44]. However, this by itself does not explain growth arrest, as body growth in larvae is primarily driven by cell volume growth, rather than cell proliferation [45]. On the other hand, growth arrest does not simply reflect a lack of food: it is observed not only for starvation, but also for stresses such as heat or osmotic shock, where in principle sufficient food is present to support growth [23,24]. Therefore, we examined if body growth itself was also actively regulated by IIS and DAF-16/FOX.

As measure of body growth, we quantified the length of animals along their A-P axis as function of time, for unstressed and different stressed conditions. In the absence of stress, body length increased approximately linearly over the course of the L1 larval stage, as described previously [27,46,47], but such extension was absent upon starvation (**SI Fig. 5A,B**). While body growth was similarly arrested for 300 mM NaCl osmotic stress (**SI Fig. 5C**), we observed a surprising variability in growth arrest for 200 mM NaCl (**Fig. 4A**), with some animals showing fully linear growth, albeit at lower rate than without stress, while others displayed complete growth arrest or intermediate growth. Strikingly, we found that the amount of growth depended on the number and duration of DAF-16::GFP pulses (**Fig. 4A, SI Fig. 5D**). Animals with approximately normal growth typically showed few or no DAF-16::GFP pulses, while fully arrested animals showed persistent DAF-16::GFP pulse trains. Moreover, especially for intermediate growth, individual pulses often coincided with a temporary halt in growth. This indicated that DAF-16::GFP translocation itself was directly linked to body growth.

**Figure 4.**
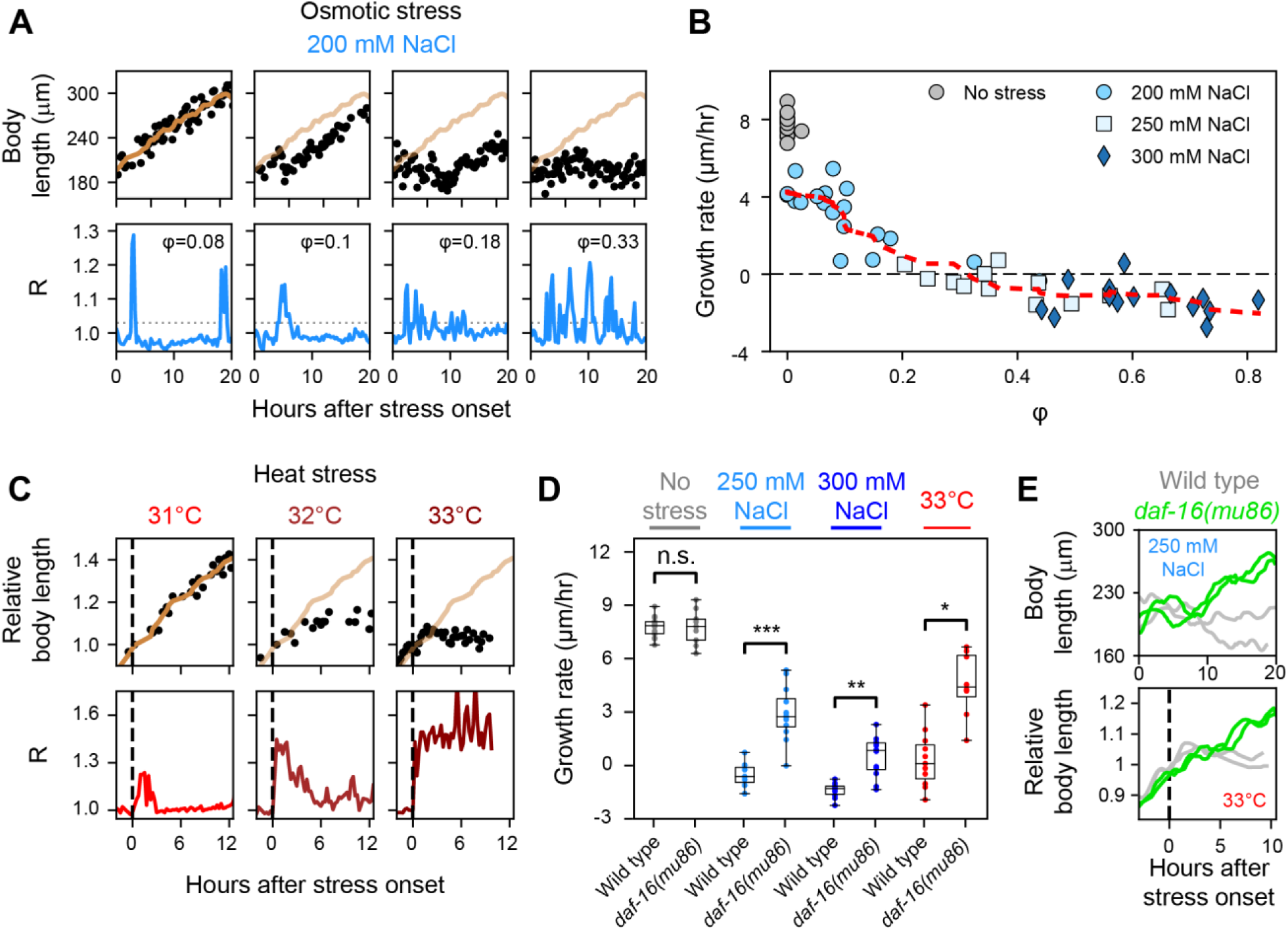
DAF-16/FOXO translocation pulses cause arrest of body growth. **(A)** Body length (top) and DAF-16::GFP nuclear localization (*R*, bottom) for individual animals under 200 mM NaCl osmotic shock, for increasing fractions of time ϕ that DAF-16::GFP was nuclear. Black markers are individual measurements. Orange line is length data for the animal with ϕ =0.08 smoothened with a S-G filter, shown for comparison. Sustained pulses or trains of shorter pulses correspond to times of reduced body growth. **(B)** Growth rate as function of *ϕ* for different magnitude of osmotic shock. Growth rate is defined as Δ*L/*Δ*T*, where Δ*L* is the change in body length over the measured time interval Δ*T*. Each marker represents an individual animal. Red line is smoothed growth rate data (S-G filter). The apparent negative growth rate for *ϕ* >0.3 is due to osmotic-shock driven decrease in body volume that is independent of growth. **(C)** Same as (A) but for individuals under increasing heat shock. Dashed line is time of temperature shift. Body length is measured relative to length at time of heat shock, which in each individual occurs at a different time relative to hatching. **(D)** Boxplots comparing growth rate for wild-type and *daf-16(mu86)* animals under various stress conditions. *daf-16(mu86)* mutants sustain growth under all stress conditions, meaning that DAF-16 is required for reduced or arrested growth under osmotic stress or heat shock. *:P<1·10^−3^, **:P<1·10^−4^, ***:P<1·10^−5^, N.S.: not significant (Welch’s t-test). (E) Body length for two wild-type (grey) and two *daf-16(mu86)* animals exposed to 250 mM NaCl (top) and 33°C (bottom). Length data was smoothened as in (A).

To further examine the relation between DAF-16::GFP pulses and growth, we quantified each animal’s average growth rate and compared it to the fraction of time ϕ that DAF-16::GFP was nuclear (defined as *R*>1.03), for animals exposed to 200, 250 and 300 mM NaCl (**Fig. 4B**). The measured growth rate for animals under osmotic stress was always lower than for non-stressed animals, even when no DAF-16::GFP pulses were observed, and was negative for animals in 300 mM NaCl. Both observations likely reflected the known reduction in body volume induced by osmotic shock [48] that is independent of growth arrest. For frequent DAF-16::GFP pulses (ϕ>0.2), we found that animals ceased growth completely, consistent with growth arrest. However, for few DAF-16::GFP pulses (ϕ<0.2), most animals showed persistent growth, with a rate that was largely independent of ϕ. Variability in growth arrest and intermediate growth phenotypes were seen at the border between these two regimes. Interestingly, while ϕ varied significantly between animals under the same osmotic stress conditions, the measured growth rates for animals under all osmotic stress conditions appeared to fall onto a single curve that only depended on ϕ, indicating that each animal’s ϕ is a better predictor of growth rate than external stress conditions per se. Overall, these observations imply that animals integrate over DAF-16::GFP pulse dynamics, meaning that growth arrest is only enacted when the animal experiences a sufficient number of pulses, or pulses with sufficiently long duration, within a certain interval of time. We observed similar patterns, with some differences, for heat shock (**Fig. 4C**). In contrast to osmotic shock, most animals at 31°C initially grew with a rate similar to unstressed animals, despite DAF-16::GFP exhibiting its first, transient translocation pulse. However, pulses in DAF-16::GFP nuclear enrichment occurring after ∼5 hr typically coincided directly with cessation of growth (**Fig. 4C**, **SI Fig. 5E**). Consistently, the earlier occurrence of such nuclear enrichment pulses seen for animals on 32°C was accompanied by an earlier growth arrest. Finally, for 33°C, when DAF-16::GFP showed high nuclear enrichment in all animals, we observed full growth arrest immediately after onset of heat stress (**Fig. 4C**).

The striking coincidence of DAF-16::GFP nuclear translocation pulses and (transient) halt in growth (**Fig. 4A**, ϕ=0.1 and 0.18) strongly suggested that DAF-16 nuclear localization directly drove growth arrest. However, it could also be explained by an alternative hypothesis, namely that DAF-16::GFP nuclear translocation and growth arrest were controlled independently and in parallel by an upstream stress signal. To differentiate between these two scenarios, we placed animals carrying a *daf-16(mu86)* null allele under the same stress conditions. We found that *daf-16(mu86)* animals exposed to osmotic shock (250 or 300 mM NaCl) or heat shock (33°C) showed significantly higher growth than wild-type animals, which typically showed full growth arrest under these conditions (**Fig. 4D, E**). These results indicate that stress-induced growth arrest requires DAF-16 function and, thus, that the DAF-16 translocation pulses cause the growth arrest.

### FOXO3A nuclear translocation pulses in human cells

Stochastic translocation pulses had so far not been observed for FOXO TFs in general, yet our findings suggest that such pulses are an inherent feature of insulin signaling. To investigate whether this is conserved across animals, we turned to human cell culture. DAF-16 has four homologs in humans, *FOXO1*, *FOXO3A*, *FOXO4* and *FOXO6.* Among these, FOXO3 is associated with longevity [2], and has been proposed to retain ancestral functions [1]. We therefore transfected the human osteosarcoma U2OS cell line with a plasmid carrying FOXO3A::GFP (**Methods**). We subjected the culture to serum starvation, a condition that activates FOXO3A [8], by transferring cells from 10 % FBS to 0 % FBS, and followed FOXO3A::GFP subcellular localization for ∼20 hours. Indeed, we observed that in many cells FOXO3A translocated between the nucleus and cytoplasm in repeated pulses **(Fig. 5A,B**). These pulses occurred at a slower pace than in *C. elegans*, with a ∼5-10 h period. Unlike in the *C. elegans* experiments, which did not permit titration of the starvation signal, the media concentration of FBS could be systematically adjusted. This allowed us to investigate whether pulse dynamics depended on stress magnitude, by placing cells in 1% and 0.1% FBS. Indeed, we observed that 0.1% FBS reduced the amplitude of the pulses, and 1% FBS resulted in mostly cytoplasmic localization (**Fig. 5C**). These results suggest that, depending on the concentration of growth factors and nutrients, the pulses can be modulated to control the time that FOXO3A spends in the nucleus. Overall, our results indicate that pulsatile regulation by FOXO transcription factors might be a general feature of the insulin signaling pathway.

**Figure 5.**
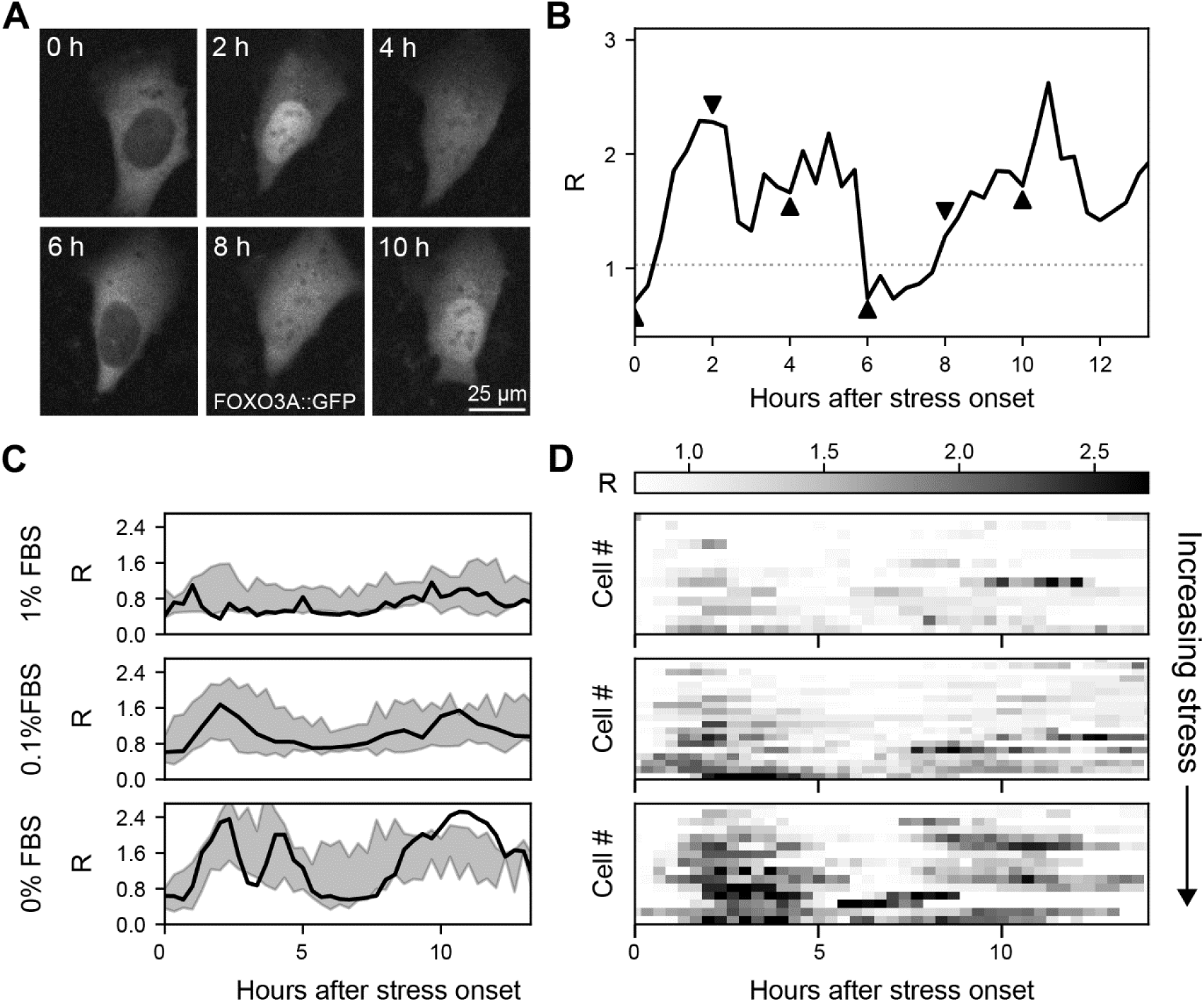
Stress-induced FOXO3A nuclear translocation pulses in human cells. **(A)** Fluorescence images of FOXO3A::GFP in a single U2OS cell under nutrient starvation (0% FBS in the medium), at different times since the start of the experiment. Scale bar: 25 µm. **(B)** FOXO3A::GFP nuclear localization *R* for the cell imaged in (A), with arrowheads indicating the time of each image. *R* =1 corresponds to equal FOXO3A::GFP intensity in nucleus and cytoplasm. **(C)** Individual cell tracks and (D) population-view heatmaps (right) of *R*, for increasing nutrient starvation (decreasing % FBS). Individual tracks are compared with the standard deviation for the cell population (shaded regions).

## Discussion

FOXO TFs, including DAF-16/FOXO in *C. elegans*, are currently assumed to remain cytoplasmic under favorable conditions and become nuclearly localized upon unfavorable conditions, where they induce genes required for stress resistance and inhibit genes for proliferation and growth [7,22,36,49,50]. Here, our dynamic measurements of DAF-16/FOXO translocation dynamics during stress-induced L1 arrest revealed a strikingly different picture. We found that under constant stress (starvation, osmotic shock or heat), DAF-16/FOXO entered and exited the nucleus in distinct pulses that were highly synchronized between the different cells in the body (**Figs. 1,2**). While DAF-16/FOXO nuclear localization increased with stress magnitude, DAF-16/FOXO pulse dynamics differed qualitatively between different types of stress at all stress magnitudes examined (**Fig. 2**). We observed similar nuclear translocation pulses of FOXO3A in a human cell line upon nutrient stress (**Fig. 5**), indicating that such pulses are a general feature of FOXO TFs. In general, these result show that studying FOXO translocation by examining single microscopy snapshots is insufficient and that, instead, dynamic measurements are essential.

A surprising aspect of DAF-16/FOXO translocation pulses is that they are inherently stochastic, with the incidence of an individual pulse typically difficult to predict, yet occur simultaneously in all *daf-16* expressing cells throughout the body. Our results identified a function for such synchronized pulses, namely in coordinating body-wide growth. This is based on the close relationship observed between DAF-16/FOXO pulse dynamics and growth arrest (**Fig. 4**), especially for animals under osmotic stress: for short (< 0.5 h), isolated pulses animals did not show any growth arrest, while sufficiently persistent DAF-16/FOXO pulse trains resulted in complete lack of growth. This was further supported by our observation of intermediate cases, where an isolated DAF-16/FOXO pulse of longer duration (> 0.5 hr) caused an immediate, but transient growth arrest. Overall, these results suggest that cells integrated over DAF-16/FOXO pulses, so that growth arrest was only induced upon sustained DAF-16/FOXO nuclear localization and not induced by transient translocation pulses. This observed link between DAF-16/FOXO pulse dynamics and body-wide growth provides a compelling rationale for the observed synchrony: to maintain uniform proportions, (arrest of) cell volume growth and, hence, pulse dynamics, must be tightly synchronized between all cells.

The lack of observed growth arrest in *daf-16* loss-of-function mutants under osmotic stress and heat shock (**Fig. 4**) indicated that cessation of growth was a direct consequence of DAF-16/FOXO nuclear translocation. While the role of DAF-16/FOXO in *C. elegans* developmental arrest is well-established, it was so far only linked to arrest of the cell cycle [22,44] rather than of cell volume increase, which is the main driver of body growth [45]. Our finding that DAF-16/FOXO controls body-wide growth of *C. elegans* larvae is therefore novel. It also matches earlier results that *Drosophila* IIS regulates organ and cell size [13,51–54]. These previous studies left unaddressed how FOXO proteins control size. One proposed mechanism is that IIS impacts cell size through direct control of protein synthesis, by impacting proteins, such as initiation factors or ribosomal proteins, that set translation rates [53,54]. In the alternative mechanism, growth is controlled more indirectly, by FOXO inducing large-scale changes in gene expression, including of metabolic genes, and the resulting rerouting of metabolism impacting growth [43,55]. Our work now provides the first examination of IIS activity and body-wide growth at high time resolution. Strikingly, we observed several cases where DAF-16/FOXO nuclear translocation was rapidly (within 0.5 h) followed by full arrest in body length extension, in particular for osmotic shock (**Fig. 4A, SI Fig. 5D**). In contrast, full induction of L1 arrest-specific gene expression, including many metabolic genes, was shown to require 3-6 h [43,56], too long to explain the rapid impact on growth we observed. Our results thus favor a mechanism where IIS controls growth directly, on timescales short compared to expression of metabolic genes.

Individual FOXO family members respond to multiple stresses, by inducing stress-specific gene-expression programs [7–9,49]. Indeed, in *C. elegans* the majority of genes expressed under osmotic shock were not induced under starvation [23], even though in both cases stress response is mediated by IIS and DAF-16/FOXO nuclear translocation. How a single signaling pathway, such as IIS, can respond to different stresses with a tailored gene expression response is an important, unresolved question. In general, it has been speculated that stress-specificity of FOXO-induced gene expression is achieved by interaction with other TFs [29,49,57,58]. In *C. elegans,* for example, the TFs HSF-1 and HCF-1 were proposed to direct DAF-16/FOXO to specifically induce expression of either heat-shock (HSF-1) or oxidative stress (HCF-1) response genes [59,60]. Our observation that DAF-16/FOXO translocation dynamics differed qualitatively between stress types (**Fig. 2**), ranging from stochastic oscillations (starvation) to random pulses (osmotic shock) or a single pulse of fixed duration (heat shock), now suggests an alternative mechanism for stress discrimination. Given that the ability of DAF-16/FOXO to induce gene expression depends crucially on its nuclear localization, differences in the duration or frequency of DAF-16/FOXO translocation between stresses will likely impact gene expression. Indeed, while novel for FOXO TFs, pulsatile translocation dynamics has been discovered in recent years more broadly in signaling networks linked to stress response [61,62], and single-cell studies have shown that stress-specific differences in TF nuclear translocation dynamics are sufficient to induce expression of different genes [62,63]. For example, transient and sustained nuclear TF localization often leads to expression of distinct sets of genes [64,65], while changes in number and duration of TF translocation pulses can cause expression of different genes, depending on the nature of their promoters [66]. Hence, we speculate that the observed stress-specificity of translocation dynamics are key in explaining how a single TF, in this case DAF-16/FOXO, induces a stress response that is specific to stress type.

While pulses in nuclear translocation dynamics have been observed for pathways, such as Erk and P53, that govern developmental processes and stress response in animals [65,67], they have so far only been examined in isolated cells. Here, we used *C. elegans* to study, for the first time, TF translocation pulses on the cellular level, but within the context of the animal’s body. Our observation that DAF-16/FOXO translocation pulses were variable between individual animals, yet strongly synchronized throughout the animal’s body (**Figs. 1,2**) implies the action of a communication mechanism that maintains strong synchrony between cells long after the shift to stressed conditions. Our measurements indicate that this communication must be fast, as delays in DAF-16/FOXO translocation between intestinal cells were of order 1-3 min (**Fig. 2**), much shorter than the ∼20 min required for DAF-16/FOXO to transition from cytoplasmic to nuclear, or vice versa. Our data for starvation revealed that DAF-16/FOXO translocation occurred slightly earlier in anterior versus posterior intestinal cells, suggestive of a head-to-tail signal responsible for synchronization and consistent with proposed long-range signals emanating from head neurons that detect starvation [68]. However, synchronization could arise without long-range signals purely through local coupling between cells, as was demonstrated for example for the collective, wave-like gene expression oscillations in somitogenesis [69].

A compelling open question concerns the molecular identity of the synchronizing signal. An intriguing candidate are the insulin-like peptides (ILPs). While *C. elegans* has only a single insulin-like receptor, DAF-2, its genome encodes 40 ILPs that function as either agonists or antagonists of IIS [70]. ILPs are excreted by multiple tissues, including neurons, intestine and hypodermis [29]. Moreover, it was shown that DAF-16/FOXO activation by IIS in one tissue induced ILP expression that in turn impacted IIS in other tissues [71,72]. In principle such inter-tissue IIS interactions appear sufficiently complex to synchronize and possibly even generate DAF-16/FOXO translocation pulses, through body-wide modulation of ILP secretion. However, IIS-mediated changes in ILP expression were shown to require 1-6 h [71], which is relatively slow compared to the observed timescale of DAF-16/FOXO translocation pulses (<1 h). It will therefore be interesting to examine whether IIS can impact ILP excretion also on shorter timescales.

The molecular machinery of IIS is maintained from nematodes to mammals, with a conserved role in organizing body-wide responses. Our unexpected observation of nuclear translocation pulses both for DAF-16/FOXO in *C. elegans* and for FOXO3A in human cell lines indicate that such pulses are likely a broadly conserved feature of FOXO signaling. It will be important to establish whether FOXO translocation pulses, including long-range synchronization to ensure body-wide coordination, also function in higher organisms.

## Acknowledgements

We thank Peter Askjaer, Tom Shimizu and Pieter Rein ten Wolde for comments and critical Reading of the manuscript. Initial observations of pulses were performed at the lab of J.v.Z during a COST GENiE Short Term Scientific Misssion granted to M.O. (COST-STSM-BM1408-35915). O.F., Y.G. and J.v.Z. were funded by the VIDI grant 680-47-529 financed by the Dutch Research Council (NWO). T.L was funded by the project OCENW.KLEIN.072 financed by the Dutch Research Council (NWO). Work in the Olmedo lab was supported by the grant PID2019-104632GB-I00 funded by MCIN /AEI/10.13039/501100011033. M.A.S-R. was supported by the VI Reseach Plan of the University of Sevilla (PPIT-US). Some strains were provided by the Caenorhabditis Genetics Center (CGC), which is supported by the National Institutes of Health-Office of Research Infrastructure Programs (P40 OD010440).

## Author contributions

O.F., B.D., M.O. and J.v.Z. conceived the research. O.F., B.D., M.O. and J.v.Z. wrote the manuscript with the input and discussion of all authors. O.F., B.D., T.L, Y.G and M.O. performed *C. elegans* experiments. M.A.S-R. performed cell culture experiments. O.F., B.D. and T.L. performed analysis of microscopy data. O.F. and B.D. developed Python scripts to perform most data analysis.

---

**SI Figure S1.**
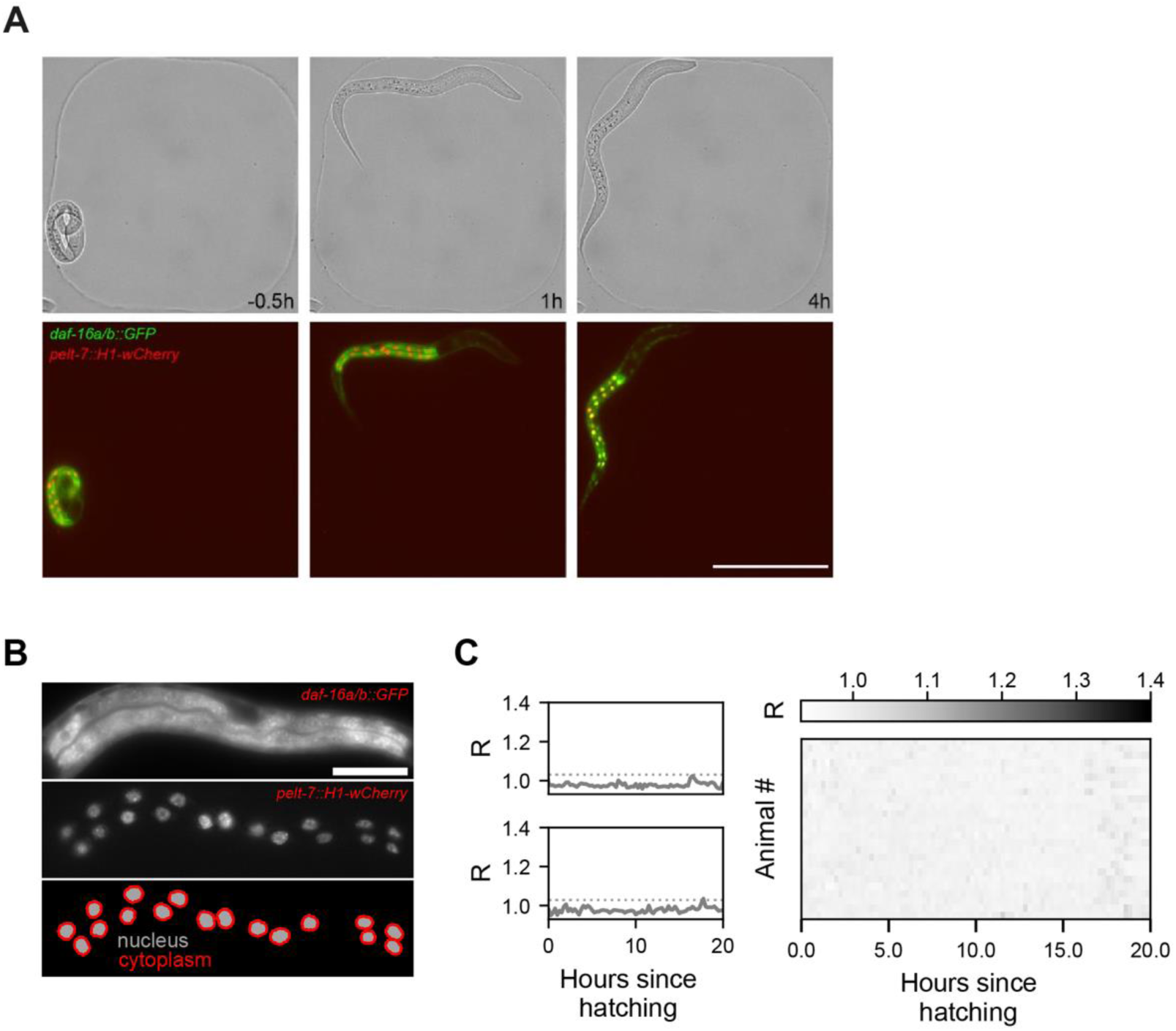
Automated quantification of DAF-16 pulse dynamics. **(A)** Single L1 larva imaged in a microchamber. Time is hours relative to hatching. Scale bar: 100 μm. **(B)** Fluorescence images of DAF-16::GFP (top panel) and intestinal marker H1-wCherry (middle panel) in the intestinal cells of an L1 larva. Bottom panel shows the binary mask of the nuclei (gray) and cytoplasm (red) of the intestinal cells for which the DAF-16::GFP fluorescence intensity was calculated. Note that the cytoplasmic mask only covers a narrow strip of cytoplasm directly surrounding each nucleus. Scale bar: 20μm. **(C)** Individual tracks (left) and population view heatmaps (right) of DAF-16::GFP nuclear localization *R* averaged over all intestinal cells, for animals on food. DAF-16::GFP remains cytoplasmic at almost all times, as expected.

**SI Figure S2.**
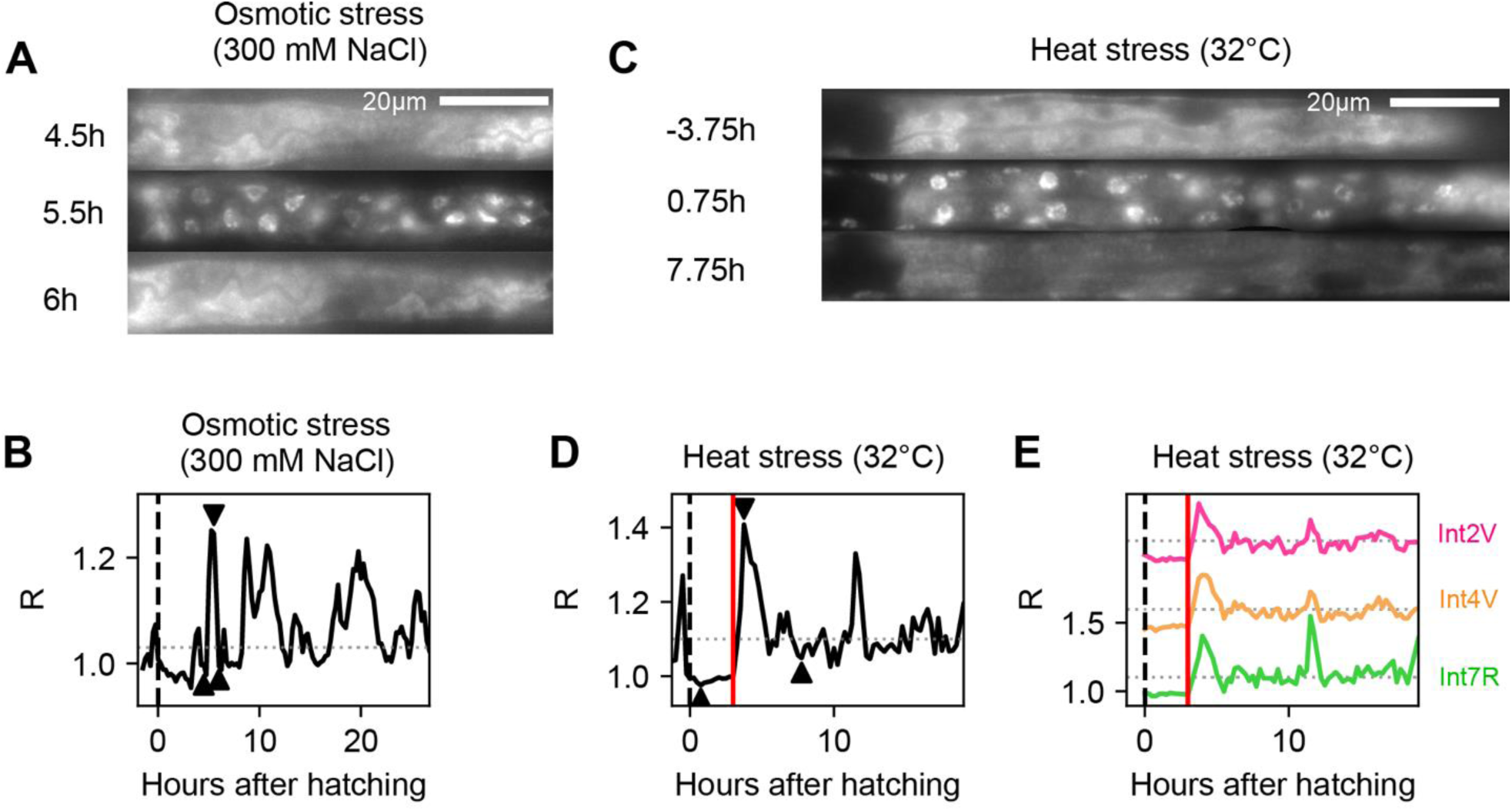
DAF-16 dynamics under osmotic and heat stress. **(A)** Fluorescence images of DAF-16::GFP in the intestinal cells of an L1 larva subject to osmotic stress (300 mM NaCl). Time is hours since the onset of osmotic shock at the time of hatching. Images were computationally straightened for clarity. Scale bar: 20 µm. **(B)** DAF-16::GFP nuclear localization *R* of the animal in (A), averaged over all intestinal cells. Arrowheads correspond to the time points in (A). **(C)** Fluorescence images of DAF-16::GFP in the intestinal cells of an L1 larva under 32°C heat shock. Time in hours relative to the onset of heat stress. Animals under heat shock are larger that animals under osmotic shock both because the former entered L1 arrest at the time of the temperature shift, rather than immediately upon hatching, and because animals shrink under osmotic shock. Scale bar: 20 µm. **(D)** DAF-16::GFP localization trajectory *R* for the animal in (C), averaged over all intestinal cells. Arrowheads correspond to the time points shown in (C), with the red line indicating the time of shift from 20°C to 32°C. **(E)** Same animal as (C) and (D), but here *R* is shown for individual intestinal cells. Trajectories are shifted for clarity.

**SI Figure S3.**
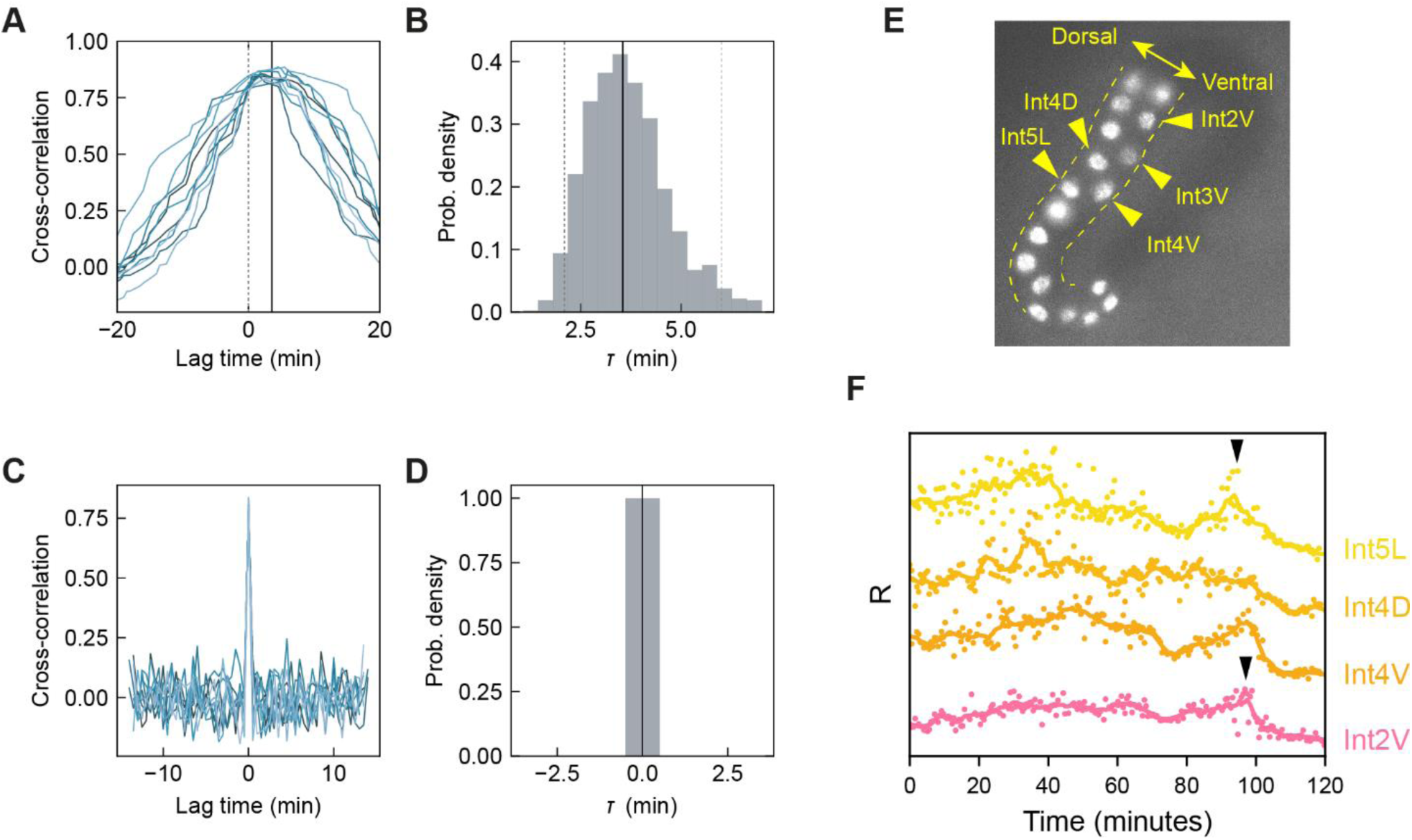
DAF-16::GFP translocation delay between intestinal cells. **(A)** Cross-correlations of DAF-16::GFP localization trajectories between Int2V and Int8R cells. The 10 different curves are generated from 10 different subsets randomly sampled from a single worm using Monte Carlo simulations. The average peak position or delay (τ, solid black line) clearly differs from 0 (dashed gray line). **(B)** Distribution of peak positions or delays for 1000 cross-correlation random subsets. **(C)** Cross-correlations of scrambled DAF-16 localization trajectories (see **Methods**), between Int2V and Int8R. **(D)** Distribution of peak positions or delays for 1000 cross-correlation subsets of scrambled trajectories. For the scrambled data there is no delay, which indicates that the delay observed between cells with unshuffled data is not an artifact. **(E)** Example of an animal outlining dorsal location of Int4D, 5L and ventral location of Int2V, Int3V and Int4V. Fluorescence is an intestinal nuclear marker. Dashed lines indicate the relevant part of the body outline. **(F)** Example trajectories of nuclear localization *R* showing dorsal-ventral order in DAF-16::GFP translocation dynamics upon osmotic shock. Black arrow marks an example of a DAF-16::GFP nuclear translocation pulse that occurs in Int5L before Int2V.

**SI Figure S4.**
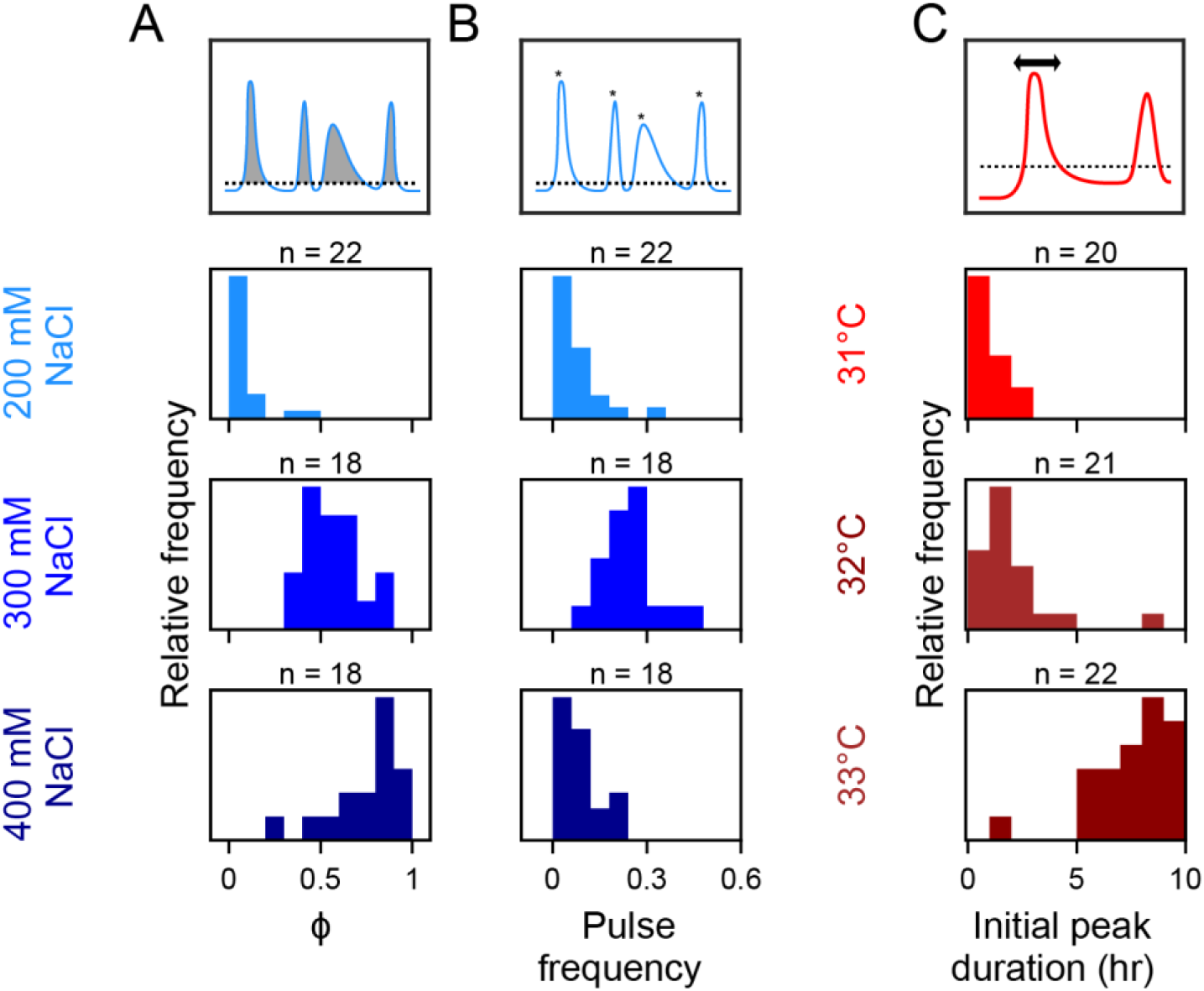
Quantification of pulse characteristics under osmotic and heat stress. **(A), (B)** Distributions of (A) the fraction of time ϕ that DAF-16::GFP is nuclear and (B) pulse frequency, both for increasing osmotic shock. **(C)** Distributions of peak duration of the first DAF-16::GFP translocation peak for increasing heat shock.

**SI Figure S5.**
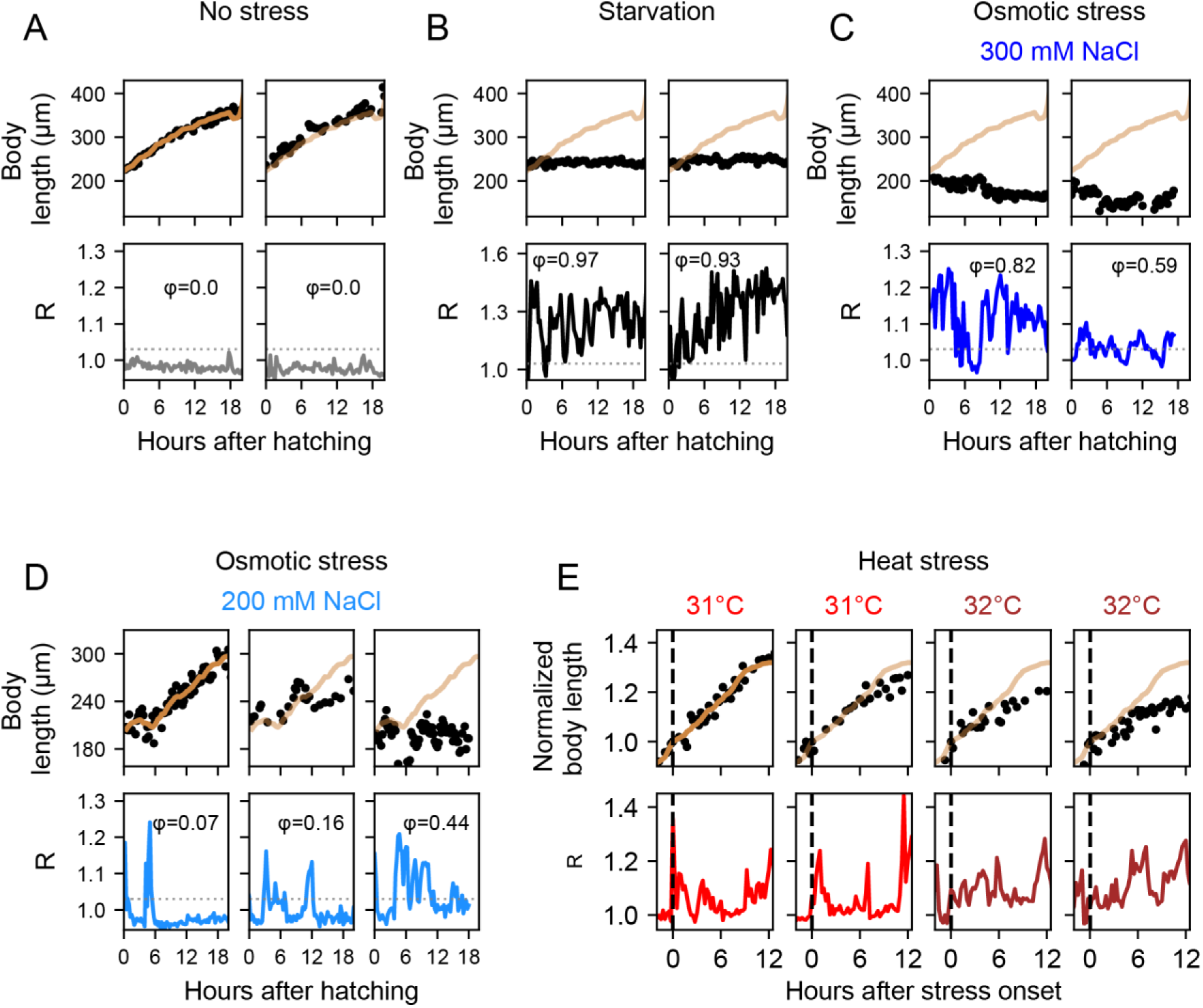
Growth arrest coincides with DAF-16 pulses. **(A)** Body length (top) and DAF-16::GFP nuclear localization (*R*, bottom) for two individuals in absence of stress. Animals grow at a constant rate, which decreases as they approach ecdysis. The smoothened length data (orange, Savitzky-Golay filter) of the animal in the left panel is also shown in the right panel for comparison. **(B),(C)** Same as (A), but under starvation (B) or osmotic stress (C). For starvation, animals do not show any growth. Under 300 mM NaCl osmotic shock, most animals at the end of imaging have not noticeably grown, and in many cases even shrunk. Orange line is length data for the unstressed animal in (A), shown here for comparison. **(D)** Individuals under 200 mM NaCl osmotic shock. Growth rate is linked to number and duration of DAF-16 pulses, with timing of growth arrest coinciding with DAF-16 pulses. Orange line is smoothened length data of the animal with *ϕ*=0.07 and sustained growth, shown in other panels for comparison. **(E)** Individuals exposed to different heat stresses. Upon heat shock, animals do not arrest growth after the first DAF-16::GFP pulse, but only after secondary pulse(s). Orange line is smoothened length data of the animal with sustained growth in the left panel, shown in other panels for comparison.

## References

1. Gui, T. and B.M.T. Burgering, FOXOs: masters of the equilibrium. Febs j, 2022. 289(24): p. 7918–7939.

2. Willcox, B.J., et al., FOXO3A genotype is strongly associated with human longevity. Proc Natl Acad Sci U S A, 2008. 105(37): p. 13987–92.

3. Flachsbart, F., et al., Identification and characterization of two functional variants in the human longevity gene FOXO3. Nat Commun, 2017. 8(1): p. 2063.

4. Gross, D.N., M. Wan, and M.J. Birnbaum, The role of FOXO in the regulation of metabolism. Curr Diab Rep, 2009. 9(3): p. 208–14.

5. Jiramongkol, Y. and E.W. Lam, FOXO transcription factor family in cancer and metastasis. Cancer Metastasis Rev, 2020. 39(3): p. 681–709.

6. Wang, Y., Y. Zhou, and D.T. Graves, FOXO transcription factors: their clinical significance and regulation. Biomed Res Int, 2014. 2014: p. 925350.

7. Kops, G.J., et al., Forkhead transcription factor FOXO3a protects quiescent cells from oxidative stress. Nature, 2002. 419(6904): p. 316–21.

8. Nemoto, S., M.M. Fergusson, and T. Finkel, Nutrient availability regulates SIRT1 through a forkhead-dependent pathway. Science, 2004. 306(5704): p. 2105–8.

9. Tran, H., et al., DNA repair pathway stimulated by the forkhead transcription factor FOXO3a through the Gadd45 protein. Science, 2002. 296(5567): p. 530–4.

10. Brunet, A., et al., Akt promotes cell survival by phosphorylating and inhibiting a Forkhead transcription factor. Cell, 1999. 96(6): p. 857–68.

11. Puig, O., et al., Control of cell number by Drosophila FOXO: downstream and feedback regulation of the insulin receptor pathway. Genes Dev, 2003. 17(16): p. 2006–20.

12. Baker, J., et al., Role of insulin-like growth factors in embryonic and postnatal growth. Cell, 1993. 75(1): p. 73–82.

13. Brogiolo, W., et al., An evolutionarily conserved function of the Drosophila insulin receptor and insulin-like peptides in growth control. Curr Biol, 2001. 11(4): p. 213–21.

14. Liu, J.P., et al., Mice carrying null mutations of the genes encoding insulin-like growth factor I (Igf-1) and type 1 IGF receptor (Igf1r). Cell, 1993. 75(1): p. 59–72.

15. Shingleton, A.W., et al., The temporal requirements for insulin signaling during development in Drosophila. PLoS Biol, 2005. 3(9): p. e289.

16. Tennessen, J.M. and C.S. Thummel, Coordinating growth and maturation - insights from Drosophila. Curr Biol, 2011. 21(18): p. R750–7.

17. Petersen, M.C. and G.I. Shulman, Mechanisms of Insulin Action and Insulin Resistance. Physiol Rev, 2018. 98(4): p. 2133–2223.

18. Chávez, V., et al., Oxidative stress enzymes are required for DAF-16-mediated immunity due to generation of reactive oxygen species by Caenorhabditis elegans. Genetics, 2007. 176(3): p. 1567–77.

19. Henderson, S.T., M. Bonafè, and T.E. Johnson, daf-16 protects the nematode Caenorhabditis elegans during food deprivation. J Gerontol A Biol Sci Med Sci, 2006. 61(5): p. 444–60.

20. Ogg, S., et al., The Fork head transcription factor DAF-16 transduces insulin-like metabolic and longevity signals in C. elegans. Nature, 1997. 389(6654): p. 994–9.

21. Weinkove, D., et al., Long-term starvation and ageing induce AGE-1/PI 3-kinase-dependent translocation of DAF-16/FOXO to the cytoplasm. BMC Biol, 2006. 4: p. 1.

22. Baugh, L.R. and P.W. Sternberg, DAF-16/FOXO regulates transcription of cki-1/Cip/Kip and repression of lin-4 during C. elegans L1 arrest. Curr Biol, 2006. 16(8): p. 780–5.

23. Burton, N.O., et al., Insulin-like signalling to the maternal germline controls progeny response to osmotic stress. Nat Cell Biol, 2017. 19(3): p. 252–257.

24. Muñoz, M.J. and D.L. Riddle, Positive selection of Caenorhabditis elegans mutants with increased stress resistance and longevity. Genetics, 2003. 163(1): p. 171–80.

25. Kaplan, R.E.W., et al., Food perception without ingestion leads to metabolic changes and irreversible developmental arrest in C. elegans. BMC Biol, 2018. 16(1): p. 112.

26. Mata-Cabana, A., et al., Social Chemical Communication Determines Recovery From L1 Arrest via DAF-16 Activation. Front Cell Dev Biol, 2020. 8: p. 588686.

27. Gritti, N., et al., Long-term time-lapse microscopy of C. elegans post-embryonic development. Nat Commun, 2016. 7: p. 12500.

28. Baugh, L.R., To grow or not to grow: nutritional control of development during Caenorhabditis elegans L1 arrest. Genetics, 2013. 194(3): p. 539–55.

29. Murphy, C.T. and P.J. Hu, Insulin/insulin-like growth factor signaling in C. elegans. WormBook, 2013: p. 1–43.

30. Paradis, S. and G. Ruvkun, Caenorhabditis elegans Akt/PKB transduces insulin receptor-like signals from AGE-1 PI3 kinase to the DAF-16 transcription factor. Genes Dev, 1998. 12(16): p. 2488–98.

31. Rena, G., et al., Phosphorylation of the transcription factor forkhead family member FKHR by protein kinase B. J Biol Chem, 1999. 274(24): p. 17179–83.

32. Lee, R.Y., J. Hench, and G. Ruvkun, Regulation of C. elegans DAF-16 and its human ortholog FKHRL1 by the daf-2 insulin-like signaling pathway. Curr Biol, 2001. 11(24): p. 1950–7.

33. Furuyama, T., et al., Identification of the differential distribution patterns of mRNAs and consensus binding sequences for mouse DAF-16 homologues. Biochem J, 2000. 349(Pt 2): p. 629–34.

34. Schuster, E., et al., DamID in C. elegans reveals longevity-associated targets of DAF-16/FoxO. Mol Syst Biol, 2010. 6: p. 399.

35. Tepper, R.G., et al., PQM-1 complements DAF-16 as a key transcriptional regulator of DAF-2-mediated development and longevity. Cell, 2013. 154(3): p. 676–690.

36. Henderson, S.T. and T.E. Johnson, daf-16 integrates developmental and environmental inputs to mediate aging in the nematode Caenorhabditis elegans. Curr Biol, 2001. 11(24): p. 1975–80.

37. Alam, H., et al., EAK-7 controls development and life span by regulating nuclear DAF-16/FoxO activity. Cell Metab, 2010. 12(1): p. 30–41.

38. Putker, M., et al., Redox-dependent control of FOXO/DAF-16 by transportin-1. Mol Cell, 2013. 49(4): p. 730–42.

39. Riedel, C.G., et al., DAF-16 employs the chromatin remodeller SWI/SNF to promote stress resistance and longevity. Nat Cell Biol, 2013. 15(5): p. 491–501.

40. Wolff, S., et al., SMK-1, an essential regulator of DAF-16-mediated longevity. Cell, 2006. 124(5): p. 1039–53.

41. Aghayeva, U., A. Bhattacharya, and O. Hobert, A panel of fluorophore-tagged daf-16 alleles. MicroPubl Biol, 2020. 2020.

42. Kimura, K.D., et al., daf-2, an insulin receptor-like gene that regulates longevity and diapause in Caenorhabditis elegans. Science, 1997. 277(5328): p. 942–6.

43. Hibshman, J.D., et al., daf-16/FoxO promotes gluconeogenesis and trehalose synthesis during starvation to support survival. Elife, 2017. 6.

44. Schindler, A.J., L.R. Baugh, and D.R. Sherwood, Identification of late larval stage developmental checkpoints in Caenorhabditis elegans regulated by insulin/IGF and steroid hormone signaling pathways. PLoS Genet, 2014. 10(6): p. e1004426.

45. Uppaluri, S., S.C. Weber, and C.P. Brangwynne, Hierarchical Size Scaling during Multicellular Growth and Development. Cell Rep, 2016. 17(2): p. 345–352.

46. Stojanovski, K., H. Großhans, and B.D. Towbin, Coupling of growth rate and developmental tempo reduces body size heterogeneity in C. elegans. Nat Commun, 2022. 13(1): p. 3132.

47. Uppaluri, S. and C.P. Brangwynne, A size threshold governs Caenorhabditis elegans developmental progression. Proc Biol Sci, 2015. 282(1813): p. 20151283.

48. Lamitina, S.T., et al., Adaptation of the nematode Caenorhabditis elegans to extreme osmotic stress. Am J Physiol Cell Physiol, 2004. 286(4): p. C785–91.

49. Calnan, D.R. and A. Brunet, The FoxO code. Oncogene, 2008. 27(16): p. 2276–88.

50. Huang, H. and D.J. Tindall, Dynamic FoxO transcription factors. J Cell Sci, 2007. 120(Pt 15): p. 2479–87.

51. Böhni, R., et al., Autonomous control of cell and organ size by CHICO, a Drosophila homolog of vertebrate IRS1-4. Cell, 1999. 97(7): p. 865–75.

52. Kramer, J.M., et al., Expression of Drosophila FOXO regulates growth and can phenocopy starvation. BMC Dev Biol, 2003. 3: p. 5.

53. Miron, M., et al., The translational inhibitor 4E-BP is an effector of PI(3)K/Akt signalling and cell growth in Drosophila. Nat Cell Biol, 2001. 3(6): p. 596–601.

54. Radimerski, T., et al., dS6K-regulated cell growth is dPKB/dPI(3)K-independent, but requires dPDK1. Nat Cell Biol, 2002. 4(3): p. 251–5.

55. Puigserver, P., et al., Insulin-regulated hepatic gluconeogenesis through FOXO1-PGC-1alpha interaction. Nature, 2003. 423(6939): p. 550–5.

56. Baugh, L.R., J. Demodena, and P.W. Sternberg, RNA Pol II accumulates at promoters of growth genes during developmental arrest. Science, 2009. 324(5923): p. 92–4.

57. Tissenbaum, H.A., DAF-16: FOXO in the Context of C. elegans. Curr Top Dev Biol, 2018. 127: p. 1–21.

58. Zhou, K.I., Z. Pincus, and F.J. Slack, Longevity and stress in Caenorhabditis elegans. Aging (Albany NY), 2011. 3(8): p. 733–53.

59. Hsu, A.L., C.T. Murphy, and C. Kenyon, Regulation of aging and age-related disease by DAF-16 and heat-shock factor. Science, 2003. 300(5622): p. 1142–5.

60. Li, J., et al., Caenorhabditis elegans HCF-1 functions in longevity maintenance as a DAF-16 regulator. PLoS Biol, 2008. 6(9): p. e233.

61. Levine, J.H., Y. Lin, and M.B. Elowitz, Functional roles of pulsing in genetic circuits. Science, 2013. 342(6163): p. 1193–200.

62. Purvis, J.E. and G. Lahav, Encoding and decoding cellular information through signaling dynamics. Cell, 2013. 152(5): p. 945–56.

63. Venkatachalam, V., A. Jambhekar, and G. Lahav, Reading oscillatory instructions: How cells achieve time-dependent responses to oscillating transcription factors. Curr Opin Cell Biol, 2022. 77: p. 102099.

64. Litvak, V., et al., Function of C/EBPdelta in a regulatory circuit that discriminates between transient and persistent TLR4-induced signals. Nat Immunol, 2009. 10(4): p. 437–43.

65. Murphy, L.O., et al., Molecular interpretation of ERK signal duration by immediate early gene products. Nat Cell Biol, 2002. 4(8): p. 556–64.

66. Hansen, A.S. and E.K. O’Shea, Promoter decoding of transcription factor dynamics involves a trade-off between noise and control of gene expression. Mol Syst Biol, 2013. 9: p. 704.

67. Purvis, J.E., et al., p53 dynamics control cell fate. Science, 2012. 336(6087): p. 1440–4.

68. Kang, C. and L. Avery, Systemic regulation of starvation response in Caenorhabditis elegans. Genes Dev, 2009. 23(1): p. 12–7.

69. Kageyama, R., et al., Oscillatory gene expression and somitogenesis. Wiley Interdiscip Rev Dev Biol, 2012. 1(5): p. 629–41.

70. Fernandes de Abreu, D.A., et al., An insulin-to-insulin regulatory network orchestrates phenotypic specificity in development and physiology. PLoS Genet, 2014. 10(3): p. e1004225.

71. Kaplan, R.E.W., et al., Pervasive Positive and Negative Feedback Regulation of Insulin-Like Signaling in Caenorhabditis elegans. Genetics, 2019. 211(1): p. 349–361.

72. Murphy, C.T., S.J. Lee, and C. Kenyon, Tissue entrainment by feedback regulation of insulin gene expression in the endoderm of Caenorhabditis elegans. Proc Natl Acad Sci U S A, 2007. 104(48): p. 19046–50.

